# Elevated neuronal excitability and seizure susceptibility in a mouse model of *SCN2A*-related developmental and epileptic encephalopathy

**DOI:** 10.64898/2026.08.30.748006

**Authors:** Nicole A. Hawkins, Dennis M. Echevarria-Cooper, Erin E. Corbett, Christopher H. Thompson, Diana J. Ixmatlahua-Ribera, Emma C. Thompson, Scott K. Adney, Alfred L. George, Jennifer A. Kearney

**Author notes:** Corresponding Author: Jennifer A. Kearney, Ph.D., Associate Professor of Pharmacology, Northwestern University Feinberg School of Medicine 320 East Superior St, Searle 8-510, Chicago, IL 60611. denotes equal contribution (Nicole A. Hawkins, Dennis M. Echevarria-Cooper, Erin E. Corbett).

## Abstract

Pathogenic variants in *SCN2A* cause a spectrum of neurodevelopmental disorders, including developmental and epileptic encephalopathies (DEE). The patient-associated *SCN2A*-p.E430A variant selectively shifts voltage-dependence of activation, a mechanism predicted to enhance neuronal excitability. To investigate the functional consequences of the *SCN2A*-E430A variant, we generated the novel *Scn2a^E430A^*mouse model and assessed neuronal excitability, brain activity, seizure susceptibility, and phenytoin responsiveness. Heterozygous *Scn2a^E430A^*mice retained normal *Scn2a* expression levels and did not exhibit premature lethality. Hippocampal pyramidal neurons from heterozygous *Scn2a^E430A^*mice were hyperexcitable compared to wild-type neurons. Enhanced long-term potentiation was observed in the CA1 circuit of *Scn2a*^E430A^ mice. Video-EEG data revealed recurrent epileptiform discharges and spectral abnormalities, but no spontaneous generalized seizure events. In multiple seizure-induction assays, *Scn2a^E430A^* mice had no difference in latency to first seizure signs, but exhibited faster seizure generalization and more lethality compared to wild-type mice, indicating alterations in seizure propagation and/or cessation rather than seizure initiation. Pretreatment with phenytoin improved seizure outcomes across all seizure induction methods. Together, these findings demonstrate that selective disruption of *Scn2a* activation gating is sufficient to drive neuronal hyperexcitability and network dysfunction, establishing the *Scn2a^E430A^* mouse as clinically relevant and pharmacologically tractable model of *SCN2A*-related DEE.

## 1. Introduction

Pathogenic variants in *SCN2A* exert a wide range of biophysical effects on the encoded voltage-gated sodium channel Na_V_1.2, which is expressed predominantly in the central nervous system (Ben-Shalom et al., 2017; Berecki et al., 2022; George et al., 2025; Thompson et al., 2023; Wolff et al., 2017). Correspondingly, *SCN2A* variants are associated with a spectrum of heterogenous neurodevelopmental disorders (NDDs) reflecting the complex relationship between altered channel properties and phenotypes (Ben-Shalom et al., 2017; Berecki et al., 2022; Berg et al., 2024; Crawford et al., 2021; Sanders et al., 2018; Thompson et al., 2023; Wolff et al., 2017).

In general, *SCN2A* variants with Nav1.2 gain-of-function (GoF) effects at the channel level are associated with neonatal or early infantile onset epilepsy, while loss-of-function (LoF) effects are associated with intellectual disability and/or autism spectrum disorder phenotypes with or without later onset epilepsy (Ben-Shalom et al., 2017; Sanders et al., 2012; Wolff et al., 2019; Wolff et al., 2017; Zuberi, 2022). Whether an epilepsy-associated *SCN2A* variant results in GoF or LoF effects has implications for disease progression and treatment response. GoF variants are typically associated with earlier seizure onset (≤ 1 months of age) and positive therapeutic responses to sodium channel blockers like phenytoin and carbamazepine (Adney et al., 2020; Dilena et al., 2017; Wolff et al., 2017). In contrast, variants with LoF effects are associated with later onset seizures (≥ 3 months of age) and worsening of seizures in response to sodium channel blockers (Berg et al., 2024; Wolff et al., 2017). For GoF effects, additional heterogeneity comes from variable character and degree of effects on the channel, resulting in a spectrum that ranges from self-limited familial neonatal-infantile epilepsy (SeLFNIE) to developmental and epileptic encephalopathies (DEE) (Ben-Shalom et al., 2017; Berecki et al., 2018; Misra et al., 2008; Ogiwara et al., 2009; Thompson et al., 2023; Wolff et al., 2017; Yang et al., 2022). These genotype-phenotype correlations highlight the importance of *SCN2A* variant characterization and modeling approaches that reflect the heterogeneity of *SCN2A*-related epilepsies.

Several mouse models have been developed to investigate *SCN2A* GoF variants with varying biophysical effects. The *Scn2a^Q54^* model had elevated persistent current in hippocampal neurons, juvenile-onset epilepsy, and premature lethality; however, this model was hemizygous for a transgene carrying a synthetic mutation driven by a non-native promoter (Kearney et al., 2001). The patient-associated variant *SCN2A*-p.A263V has been linked to epilepsy phenotypes of varying severity and results in elevated persistent sodium current, depolarized voltage dependence of inactivation, and altered inactivation kinetics (Berecki et al., 2022; Liao et al., 2010; Wolff et al., 2017). Homozygous *Scn2a^A263V^*knock-in mice exhibited frequent, life-long spontaneous seizures; whereas heterozygotes, which reflect the allele dosage in affected humans, had only self-limited electrographic seizures as neonates (Reva et al., 2025; Schattling et al., 2016). In contrast, mice heterozygous for the DEE-associated variant *SCN2A-*p.R1882Q had spontaneous seizures starting at birth and juvenile lethality (Li et al., 2021). The p.R1882Q variant results in depolarized voltage-dependence, slowed inactivation, and enhanced persistent and resurgent sodium currents, leading to higher action potential (AP) firing frequency in excitatory cortical neurons (Berecki et al., 2018; Li et al., 2021; Mason et al., 2019; Thompson et al., 2023).

Additional mouse models capturing distinct biophysical mechanisms are needed to expand the toolkit for investigating *SCN2A-*related DEE. The missense variant *SCN2A*-p.E430A was reported as *de novo* in two unrelated individuals with early onset DEE (Feliciano et al., 2019; Wolff et al., 2017). In contrast to many GoF variants, such as p.A263V and p.R1882Q, that affect multiple channel properties, the p.E430A variant only affects voltage-dependence of activation (Thompson et al., 2023). The resulting hyperpolarizing shift in voltage-dependence of activation is predicted to increase neuronal excitability by enhancing sodium channel recruitment during subthreshold depolarizations (Berecki et al., 2018; Berecki et al., 2022; Hu and Bean, 2018; Liao et al., 2010; Liu et al., 2019; Thompson et al., 2023).

To further investigate the effects of the p.E430A variant, we generated a *Scn2a* mouse model carrying the p.E430A variant and examined effects on neuronal excitability, brain activity, seizure sensitivity, and phenytoin responsiveness. We show that *Scn2a^E430A^* hippocampal neurons have hyperpolarized action potential threshold, lower rheobase, and higher AP firing rates. Long-term potentiation in the CA1 hippocampal circuit is heightened as well, suggesting aberrant synaptic signaling, while presynaptic release probability is unaffected. At the whole animal level, *Scn2a^E430A^* mice exhibit interictal epileptiform activity and altered spectral power, as well as enhanced sensitivity to induced seizures with faster generalization and high lethality that could be attenuated by phenytoin treatment. Our results indicate that hyperpolarized Na_V_1.2 voltage-dependence of activation results in neuronal hyperexcitability, aberrant synaptic plasticity, and altered brain activity, establishing *Scn2a^E430A^* mice as a useful model of *SCN2A*-related DEE with a unique underlying mechanism.

## 2. Methods

### 2.1 Creation of structural models

To assess potential effects on structure, the protein sequence of human Na_V_1.2 (UniProt ID: Q99250-1) was used as input to AlphaFold3 with either Glu or Ala at position 430 (Abramson et al., 2024). Five structures were generated to reduce sampling bias, and models with the highest-ranking score were chosen to analyze. The Alphafold3 structure was compared with a known cryo-EM structure (PDB: 6J8E). Using PyMol, only protein regions of the AlphaFold3 structure where a corresponding cryo-EM structure was available were kept (Schrodinger, 2015). Thus, only high confidence regions (> 80% confidence) were included in analyses. Visualizations of the AlphaFold structure and cryo-EM structure were created using Chimera (Pettersen et al., 2004). To remove out-of-scope considerations, glycosylated asparagines (340, 1368, 1382, and 1393) were changed to standard asparagine, and the toxin μ-conotoxin KIIIA and a sodium ion was removed. Side chain interactions were analyzed with the Find Clashes/Contacts tool using default settings.

### 2.2 Heterologous cell electrophysiology

Heterologous expression of human Na_V_1.2 WT (Addgene #162279) E430A was performed in HEK293T cells. HEK293T cells (CRL-3216, American Type Culture Collection, Manassas, VA, USA) were maintained in Dulbecco’s modified Eagle’s medium (GIBCO/Invitrogen, San Diego, CA, USA) supplemented with 10% fetal bovine serum (Atlanta Biologicals, Norcross, GA, USA), 2 mM L-glutamine, 50 units/mL penicillin, and 50 µg/mL streptomycin at 37°C in 5% CO_2_.

The SCN2A-p.E430A variant was introduced into the full-length adult splice isoform using Q5 2X high-fidelity DNA polymerase Master Mix (New England Biolabs, Ipswich, MA) as previously described (DeKeyser et al. 2021). Whole-cell voltage-clamp recordings of voltage-gated sodium currents expressed in hetereologous cells were performed as previously described (Thompson 2011, 2012, 2020). All recordings were made at room temperature using an Axopatch 200B amplifier (Molecular Devices, LLC, Sunnyvale, CA, USA). Patch pipettes were pulled from borosilicate glass capillaries (Harvard Apparatus Ltd., Edenbridge, Kent, UK) with a multistage P-1000 Flaming-Brown micropipette puller (Sutter Instruments Co., San Rafael, CA, USA) and fire-polished using a microforge (Narashige MF-830; Tokyo, JP) to a resistance of 1.5–2.5 MΩ. The pipette solution consisted of (in mM): 10 NaF, 105 CsF, 20 CsCl, 2 EGTA, and 10 HEPES with pH adjusted to 7.35 with CsOH and osmolality adjusted to 300 mOsmol/kg with sucrose. The recording chamber was continuously perfused with bath solution containing (in mM): 145 NaCl, 4 KCl, 1.8 CaCl_2_, 1 MgCl_2_, 10 glucose and 10 HEPES with pH adjusted to 7.35 with NaOH and osmolality 310 mOsmol/kg.

### 2.3 Generation of Scn2a^E430A^ mouse model

*Scn2a^E430A^* mice (abbreviated as *Scn2a*^A/+^ or A/+) were generated using CRISPR/Cas9 editing with homology directed repair by the Northwestern University Transgenic and Targeted Mutagenesis Laboratory (TTML). Guide RNA and donor repair oligonucleotide (**Table 1**) were microinjected into C57BL/6J (B6) embryos at the two-cell stage. Potential founders were screened by Sanger sequencing of PCR products amplified using primers outside the repair oligo homology region (**Table 1**). Mosaic founders with the E430A mutation were backcrossed to B6 to generate N1 offspring. N1 offspring were genotyped by PCR and Sanger sequencing to detect transmission of the E430A editing event. Screening for off-target events included PCR and Sanger sequencing of predicted sites with < 3 mismatches (**Table 1**), and whole genome re-sequencing to survey the *Scn2a* genomic region and predicted off-targets. Heterozygous N1.*Scn2a*^A/+^ mice without detectable off-target edits were backcrossed to B6 to establish the line *Scn2a^em1Kea^* (RRID:MMRRC_075779-UCD) that is maintained as an isogenic by continual backcrossing of *Scn2a*^A/+^ heterozygotes to inbred B6 mice (B6, #000664, Jackson Laboratory). Experimental mice were generated from backcross generations ≥N3, with male and female *Scn2a*^A/+^ and WT littermates used in all experiments. At generation N10, the line was submitted to the Mutant Mouse Resource & Research Centers Repository to facilitate resource sharing (MMRRC_075779-UCD).

**Table 1.**
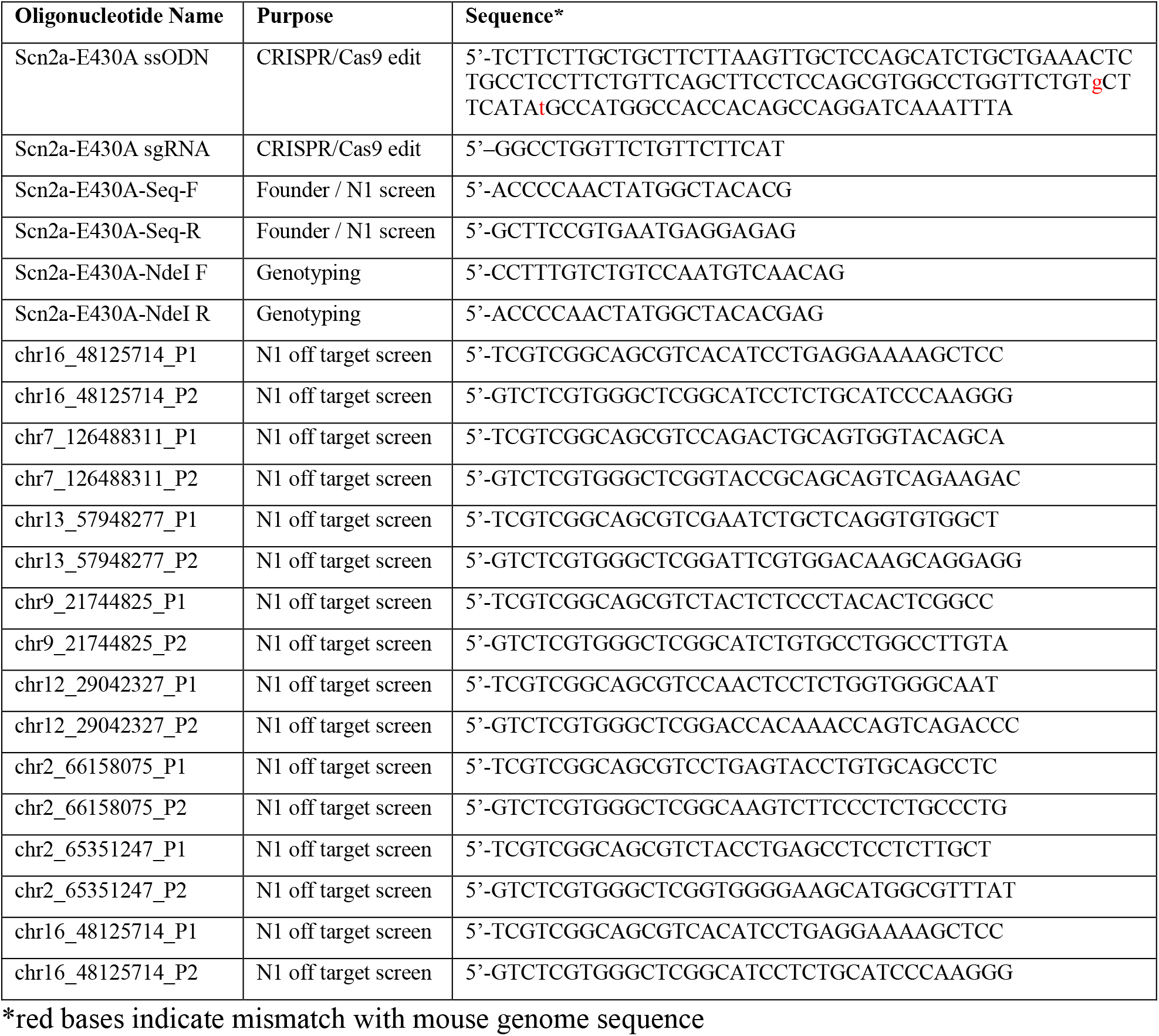
Oligonucleotides used in this study.

All mice were maintained in an SPF barrier facility with a 14:10 hour light:dark schedule with *ad libitum* access to food and water. These studies were approved by the Northwestern University Animal Care and Use Committee in accordance with the National Institutes of Health Guide for the Care and Use of Laboratory Animals. Principles outlined in the ARRIVE (Animal Research: Reporting of in vivo Experiments) guidelines were considered in planning of experiments (Percie du Sert et al., 2020).

### 2.4 Genotyping

Mice were genotyped by PCR using genomic DNA isolated from tail biopsies and a restriction fragment length polymorphism (RFLP) assay. Genomic DNA was amplified using genotyping primers (Table 1), followed by a restriction enzyme digest with Ndel (#R0111, New England Biolabs) at 37°C, ≥ 1 hour. Digestion resulted in a 539 bp fragment for WT or 539, 353, 186 bp fragments for *Scn2a*^A/+^.

### 2.5 Expression Analysis

Forebrains were isolated from 8–12-week-old male and female *Scn2a*^A/+^ and WT littermates, halved along the midline, and flash frozen and stored at-80°C until processing. For expression experiments, statistical comparisons were performed using unpaired t-test (n=4-9 per genotype).

#### 2.5.1 Transcript expression

Total RNA was isolated from one hemisphere using TRIzol (Invitrogen) and first strand cDNA was made from 4 ug of total RNA using Superscript IV Reverse Transcriptase (Thermo Fisher Scientific) according to manufacturer’s instructions. cDNA samples were diluted 1:50 and used for quantitative gene expression analysis by droplet digital PCR (ddPCR). Gene expression assays (Thermo Fisher Scientific/Applied Biosciences) targeting *Scn2a* (FAM-Mm01270359_m1) and the normalization control *Tbp* (VIC-MGB Mm00446971) were used together with ddPCR Supermix for Probes (No dUTP; Bio-Rad). Droplets were generated using a QX200 Droplet Generator (Bio-Rad) and PCR amplification was performed using standard ddPCR cycling conditions. Droplets were read on a QX200 Droplet Reader and data were analyzed using QuantaSoft v 1.7 software (Bio-Rad).

### 2.5.2 Immunoblotting

The remaining hemisphere was used for membrane protein isolation by P3 membrane protein fractionation. Protein (50 ug) was separated on a 7.5% SDS-PAGE gel and transferred to nitrocellulose membranes. Blots were probed with rabbit anti-*Scn2a* (1:200; Alomone, ASC-002) and mouse anti-mortalin/GPR75 N52A/42 (1:1000; NeuroMab/Antibodies Inc., #75-127;) primary antibodies. Alexa Fluor-conjugated secondary antibodies (α-rabbit 790 and α-mouse 680, 1:10,000; LICORbio) were used for detection on an Odyssey DLx Imaging System (LICORbio). Band intensities were quantified using Image Studio software (v. 5.2; LICORbio) and relative protein expression was calculated as the ratio of *Scn2a* to *GPR75* and normalized to WT.

### 2.6 Primary Neuron Cultures

P0–1 pups were genotyped using the RFLP genotyping assay as described above. Hippocampal neurons were harvested and plated on poly-d-lysine-coated coverslips (GG-12-1.5-PDL; Neuvitro, Vancouver WA) at a density of 0.25–0.30×10e^6^ cells per well and maintained in Neurobasal medium (10888022; Gibco, Waltham, MA) supplemented with B-27 and Culture One (17504044 and A3320201; Gibco), with weekly half-volume media changes for 2–3 weeks. At least 2 independent cultures of 2 to 3 mice of each genotype were used for experiments.

### 2.7 Neuron Electrophysiology

Whole-cell current clamp recordings were performed on DIV14–16 cultured hippocampal pyramidal neurons that were identified by their characteristic pyramidal-shaped morphology. All recordings were performed at room temperature using a MultiClamp 700B amplifier (Molecular Devices). External recording solution included (in mM): 155 NaCl, 3.5 KCl, 1.5 CaCl_2_, 1 MgCl_2_, 10 HEPES, 10 Glucose, 50 μM D-5AP (2-amino-5-phosphonopentanoic acid), 10 μM CNQX (6-cyano-7-nitroquinoxaline-2,3-dione) and 50 μM picrotoxin, with pH adjusted to 7.35 with NaOH. Internal recording solution included (in mM): 120 KMeSO_4_, 10 KCl, 5 MgATP, 0.4 Na-GTP, 5 Na_2_-phosphocreatine with pH adjusted to 7.2 with KOH, and osmolarity adjusted to 280 mOsm/kg with sucrose. Current-clamp pulse generation and data collection were done with Clampex 10.4. For evoked action potential recordings, cells were held at −65 mV and action potentials were elicited by 2 s stimuli from −30-450 pA in 10 pA increments. Input resistance was calculated from a −10 pA hyperpolarizing step. Input-output curves were compared by two-way ANOVA followed by Tukey’s post-hoc comparisons. Resting membrane potential, action potential morphology, and input resistance were compared by unpaired t-tests.

### 2.8 Slice Electrophysiology

Acute hippocampal slices were prepared from *Scn2a^A/+^* (E430A) and wild-type littermate control mice aged P27–P32. Animals were deeply anesthetized with isoflurane using a Somnosuite device (Kent Scientific Corporation) and quickly decapitated. The brain was rapidly extracted and placed in ice-cold oxygenated (95% O2/5% CO2) artificial cerebrospinal fluid (aCSF) containing in mM: 70 NaCl, 2.5 KCl, 1.25 NaH_2_PO_4_, 30 NaHCO_3_, 20 HEPES, 25 glucose, 50 sucrose, 2 thiourea, 2.5 Na L-ascorbate, 3 Na-pyruvate, 2 N-acetyl-L-cysteine, 5 Na-L-lactate, 0.5 CaCl2, 7 MgCl_2,_ 2 µM GABA, 10 µM DL-APV, 100 µM kynurenic acid, (pH 7.4, 30 mOsm/kg). Coronal 300 µm thick slices were obtained using a Leica VT1200S vibratome. Slices were allowed to recover at 32 °C for 30 minutes then were slowly changed to incubating solution containing in mM: 124 NaCl, 2.5 KCl, 1.25 NaH_2_PO_4_, 24 NaHCO_3_, 5 HEPES, 13 glucose, 2 thiourea, 2.5 Na-L-ascorbate, 3 Na-pyruvate, 2 N-acetyl-L cysteine, 5 Na-L-lacate, 1 CaCl_2_, 2 MgCl_2_, 2 µM GABA, 10 µM DL-APV, 50 µM kynurenic acid (300 mOsm/kg) oxygenated as before at room temperature. After up to 2 hours of recovery individual slices were transferred to the recording chamber and perfused with ACSF containing in mM: 124 NaCl, 2.5 KCl, 1.25 NaH_2_PO_4_, 24 NaHCO_3_, 5 HEPES, 13 glucose, 2 thiourea, 2.5 Na-L-ascorbate, 3 Na-pyruvate, 2 N-acetyl-L cysteine, 5 Na-L-lacate, 2 CaCl_2_, 1 MgCl^2^. The recording temperature was maintained at 32°C using a heating chamber connected to the TC-324C heater (Warner Instruments) at a flow rate of 2-3 mL/min. For field recordings in CA1, the slice was visualized using the SliceScope Pro 2000 (Scientifica) connected to a Kinetix 22 camera (Teledyne). The recording pipette (∼2 MΩ) was collocated on the apical dendrites of the CA1 pyramidal cells and the stimulation pipette was placed on the stratus radiatum ∼200 µm apart from the recording pipette, to stimulate the Schaffer collateral fibers on CA1 and record orthodromically-evoked field extracellular postsynaptic potentials (fEPSP). Stimulation and recording pipettes were pulled with a multi-stage P-1000 Flaming-Brown micropipette puller (Sutter Instruments Co.). Stimuli were delivered using the IsoFlex stimulator (A.M.P.I.).

The paired pulse ratio (PPR) was calculated measuring the response amplitude of two subsequent stimuli (P1 and P2) at a given interstimulus interval and compared between genotypes using two-way repeated measures ANOVA. For LTP induction, baseline evoked responses were recorded for 20 minutes, only slices with stable fEPSP amplitudes during the baseline period were included (<5% drift). Theta burst stimulation (TBS) consisted of 20 bursts of 4 pulses at 100 Hz, repeated at 5 Hz, applied at the baseline stimulation intensity. The fEPSP slope was monitored for 60 minutes post-TBS with a single stimulus delivered every 20 seconds. The LTP magnitude was quantified as the mean fEPSP slope during the 50–60-minute post-TBS window, normalized to the pre-TBS baseline and expressed as a percentage. Percentages were compared between genotypes using t-tests. Data were excluded if the series resistance changed by more than 20% during the recording or if baseline stability criteria were not met.

### 2.9 Video-EEG Recordings

WT and *Scn2a*^A/+^ mice were implanted with prefabricated 2EEG/1EMG channel headmounts (8201; Pinnacle Technology) at 4-5 months of age. Mice were anesthetized with ketamine/xylazine and reversed with Revertidine (Modern Veterinary Therapeutics). Four stainless steel screws serving as cortical surface electrodes, together with the headmount, were affixed to the skull using glass ionomer cement (GC FujiCEM). Anterior screw electrodes were positioned 0.5–1 mm anterior to bregma and 1 mm lateral from the midline, with the left anterior screw serving as the ground. Posterior screw electrodes were placed 4.5–5 mm posterior to bregma and 1 mm lateral from the midline. EEG traces correspond to recordings from the right posterior to left posterior electrodes (interelectrode distance ∼2 mm).

Following a minimum of 5 days of post-surgical recovery (range 5-18 days), tethered EEG/EMG and synchronized video data were continuously collected for ≥ 7 days using Sirenia Acquisition (Pinnacle Technology) at a sampling rate of 400 Hz. EEG recordings were segmented into lights-on (13 hours) and lights-off (9 hours) periods, excluding the 2-hour light/dark transition intervals. The first two dark segments and two light segments from each recording were manually analyzed by a reviewer blinded to genotype using LabChart v8.1.19 (ADinstruments) and Sirenia Seizure software (Pinnacle Technology). Analyses were performed using n=4 mice per genotype.

Root mean square (RMS) amplitude was calculated in LabChart for each of the four segments to represent average baseline EEG amplitude. RMS values represent two segments per mouse for each light or dark cycle and statistical comparisons were performed using unpaired t-tests. Power spectral density (PSD) was calculated from raw EEG traces using the Spectrum module in LabChart with a fast Fourier transform (FFT) size of 1024, 93.75% segment overlap, and a Hann (Cosine-bell) window applied to reduce edge effects. PSD values represent two segments per mouse for each light or dark cycle. PSD statistics were calculated by frequency band using ordinary Two-Way ANOVA with the following frequency band definitions: Delta (0-3.9 Hz) Theta (4.3-7.8 Hz) Alpha (8.2-11.7 Hz) and Beta (12.1-29.7 Hz).

### 2.10 Seizure Induction Assays

Seizure induction experiments were conducted using 6–12-week-old male and female *Scn2a*^A/+^ and WT mice, with separate cohorts for each inducer. For phenytoin (PHT) pretreatment groups, PHT (Hikma Pharmaceuticals USA, 50 mg/mL diluted in 0.5% methylcellulose) was administered 2 hours prior to seizure induction by intraperitoneal (ip) injection at 15 mg/kg. WT and *Scn2a*^A/+^ mice treated with phenytoin PHT were compared to WT and *Scn2a*^A/+^ mice with no pretreatment.

#### 2.10.1 Electroconvulsive shock (ECS)

WT and *Scn2a*^A/+^ mice were subjected to corneal electrode stimulation following topical application of 0.5% tetracaine (Sigma-Aldrich) using a 32 mA stimulus that is subthreshold for B6 mice (60 Hz, 0.5 pulse width, 0.4 sec duration; ECT unit 57800; Ugo Basile, Gemonio (VA), Italy). Within 5 seconds of stimulation, mice were assessed for the presence or absence of tonic hindlimb extension (HLE), with or without death, by an experimenter blinded to genotype. Seizure outcomes were compared by Fisher’s Exact test.

#### 2.10.2 Flurothyl. WT and Scn2a^A/+^

mice were exposed to the chemoconvulsant flurothyl (Bis(2,2,2-trifluoroethyl) ether, Sigma-Aldrich). Flurothyl was delivered into a clear plexiglass chamber (2.2 L) by a syringe pump at a constant rate of 20 μL/min and allowed to volatilize. An observer blinded to genotype recorded latency to the onset of generalized tonic-clonic seizure (GTCS) and severity classified as GTCS only, GTCS with HLE, or GTCS with HLE and death. Latencies were compared using unpaired T-tests and severity distributions were compared with Fisher’s Exact test.

#### 2.10.3 Kainic acid (KA)

Scn2a^A/+^ and WT mice were administered KA (25 mg/kg, ip; Tocris) and video monitored for 2 hours. Videos were scored offline by a reviewer blinded to genotype using the following modified Racine scale (Racine, 1972): Stage 1: Behavioral arrest; Stage 2: Forelimb extension and/or tail Straub; Stage 3: Automatisms (scratching, circling, head-bobbing); Stage 4: Forelimb clonus, rearing and/or falling; Stage 5: Repetition of stage 4; Stage 6: GTCS with loss of posture. Latency to each seizure stage was analyzed using a two-way mixed-effect ANOVA with Tukey’s *post hoc* tests. Post KA survival was also assessed, and statistical comparisons were performed using the Mantel-Cox LogRank test.

### 2.11 Statistical Analysis

Data were collected and analyzed by experimenters blinded to genotype and, when applicable, treatment group. Statistical analyses were performed using Prism v10.6.1 software (GraphPad). Initial analyses were performed separately by sex. If no sex-dependent effects were detected, data were collapsed across sex. Normality was assessed using D’Agostino & Pearson test to guide statistical test selection. All statistical tests, *post hoc* comparisons and exact p-values are reported in **Table 2** and group sizes are reported in figure legends. Data are expressed as mean ± SEM unless otherwise specified.

**Table 2.** Summary of statistical comparisons.

| Figure | Comparison | Test | Value | Post Hoc |
| --- | --- | --- | --- | --- |
| 2B | Peak Current Density | Two-way ANOVA | F(1,608)=0.1149, p=0.7347 | Tukey |
| 2C | Voltage-dependence of activation $V_{1/2}$ | Unpaired t-test | **p=0.0088** | n/a |
| 2C | Voltage-dependence of activation slope (k) | Unpaired t-test | p=0.6382 | n/a |
| 2C | Voltage-dependence of inactivation $V_{1/2}$ | Unpaired t-test | p=0.9021 | n/a |
| 2C | Voltage-dependence of inactivation slope (k) | Unpaired t-test | p=0.8077 | n/a |
| 2D | Recovery from inactivation tau fast | Unpaired t-test | p=0.8327 | n/a |
| 2D | Recovery from inactivation tau slow | Unpaired t-test | p=0.6144 | n/a |
| 2D | Recovery from inactivation percent tau fast | Unpaired t-test | p=0.9605 | n/a |
| 2E | Frequency-dependent rundown | Unpaired t-test | p=0.6178 | n/a |
|  | Persistent current | Unpaired t-test | p=0.8739 | n/a |
| 3B | Transcript | Welch's T-test | p>0.56 | n/a |
| 3C | Protein | Welch's T-test | p>0.17 | n/a |
| 4B | Input-output curves | Two-way ANOVA | **F(1,1363)=59.23 p<0.0001** | Tukey |
| 4C | RMP | Unpaired t-test | p=0.9347 | n/a |
| 4D | Rheobase | Unpaired t-test | **p<0.0001** | n/a |
| 4E | AP threshold | Unpaired t-test | **p=0.0111** | n/a |
| 4F | AP amplitude | Unpaired t-test | p=0.6705 | n/a |
| 4G | Fast AHP | Unpaired t-test | **p=0.0379** | n/a |
| 4H | AP half-width | Unpaired t-test | p=0.6398 | n/a |
| 4I | Upstroke velocity | Unpaired t-test | p=0.8988 | n/a |
| 4J | Downstroke velocity | Unpaired t-test | p=0.9162 | n/a |
|  | Input resistance | Unpaired t-test | p=0.1844 | n/a |
| 5B | Paired-pulse ratio | Two-way RM ANOVA | F(1,11)=0.2188 p=0.6491 | n/a |
| 5D | % LTP | Welch's T-test | **p=0.0288** | n/a |
| 6A | RMS (Light) | Welch's T-test | p>0.14 | n/a |
|  | RMS (Dark) | Welch's T-test | p>0.64 | n/a |
| 6B | PSD-Light (Delta) | Two-Way Anova | **F(1,154)=29.10 p<0.0001** | n/a |
|  | PSD-Light (Theta) | Two-Way Anova | **F(1,140)=4.888 p=0.0287** | n/a |
|  | PSD-Light (Alpha) | Two-Way Anova | F(1,140)=0.5385 p=0.4643 | n/a |
|  | PSD-Light (Beta) | Two-Way Anova | F(1,644)=0.1498 p=0.6989 | n/a |
| 6C | PSD-Dark (Delta) | Two-Way Anova | F(1,154)=0.5555 p=0.4572 | n/a |
|  | PSD-Dark (Theta) | Two-Way Anova | F(1,140)=3.068 p=0.0820 | n/a |
|  | PSD-Dark (Alpha) | Two-Way Anova | F(1,140)=0.8352 p=0.3623 | n/a |
|  | PSD-Dark (Beta) | Two-Way Anova | F(1,644)=0.1135 p=0.7363 | n/a |
| 7A | Electroconvulsive shock (ECS) | Fisher's Exact | **WT v A/+ p<0.0001**<br>WT PHT v A/+ PHT p>0.9999<br>**A/+ v A/+ PHT p<0.0001** | n/a |
| 7B | Flurothyl Latency to GTCS | Welch's T-test | WT v A/+ p>0.1740 | n/a |
| 7C | Flurothyl Severity | Fisher's Exact | **WT v A/+ p<0.0001**<br>**WT PHT v A/+ PHT p<0.004**<br>WT v WT PHT p=0.1954<br>**A/+ v A/+ PHT p<0.0001** | n/a |
| 7D | KA Latency to Stage<br>(Stage x Genotype) | Two-way mixed-effect ANOVA;<br>Tukey's post-hoc | **F(15,345)=6.233 p<0.0001** | *Stage 6*<br>**WT v A/+ p<0.0001**<br>**WT PHT v A/+ PHT p<0.0001**<br>**WT v WT PHT p<0.0005**<br>A/+ v A/+ PHT p>0.08 |
| 7E | KA Survival | Mantel-Cox | **WT v A/+ p<0.0001**<br>**WT PHT v A/+ PHT p<0.02**<br>WT v WT PHT p>0.06<br>**A/+ v A/+ PHT p<0.0045** | n/a |

## 3. Results

### 3.1 Effect of E430A on Na_V_1.2 channel function

Glutamate 430 is located in the proximal DI-DII cytoplasmic loop of Na_V_1.2 (**Fig. 1A**). This location is invariant across SCN2A orthologs and paralogs and is analogous to the neck region of prokaroytic voltage-gated sodium channels that has been shown to have a strong influence on voltage-dependent activation (**Fig. 1B**) (Arrigoni et al., 2016; Lenaeus et al., 2017; Sula et al., 2017). Compared to the reference sequence, Ala substitution at position 430 (E430A) predicted loss of a hydrogen bond with K247 in the DI S4-S5 linker, which is a critical region for coupling voltage-sensing with channel opening (**Fig. 1C-E**) (Catterall, 2012).

**Figure 1.**
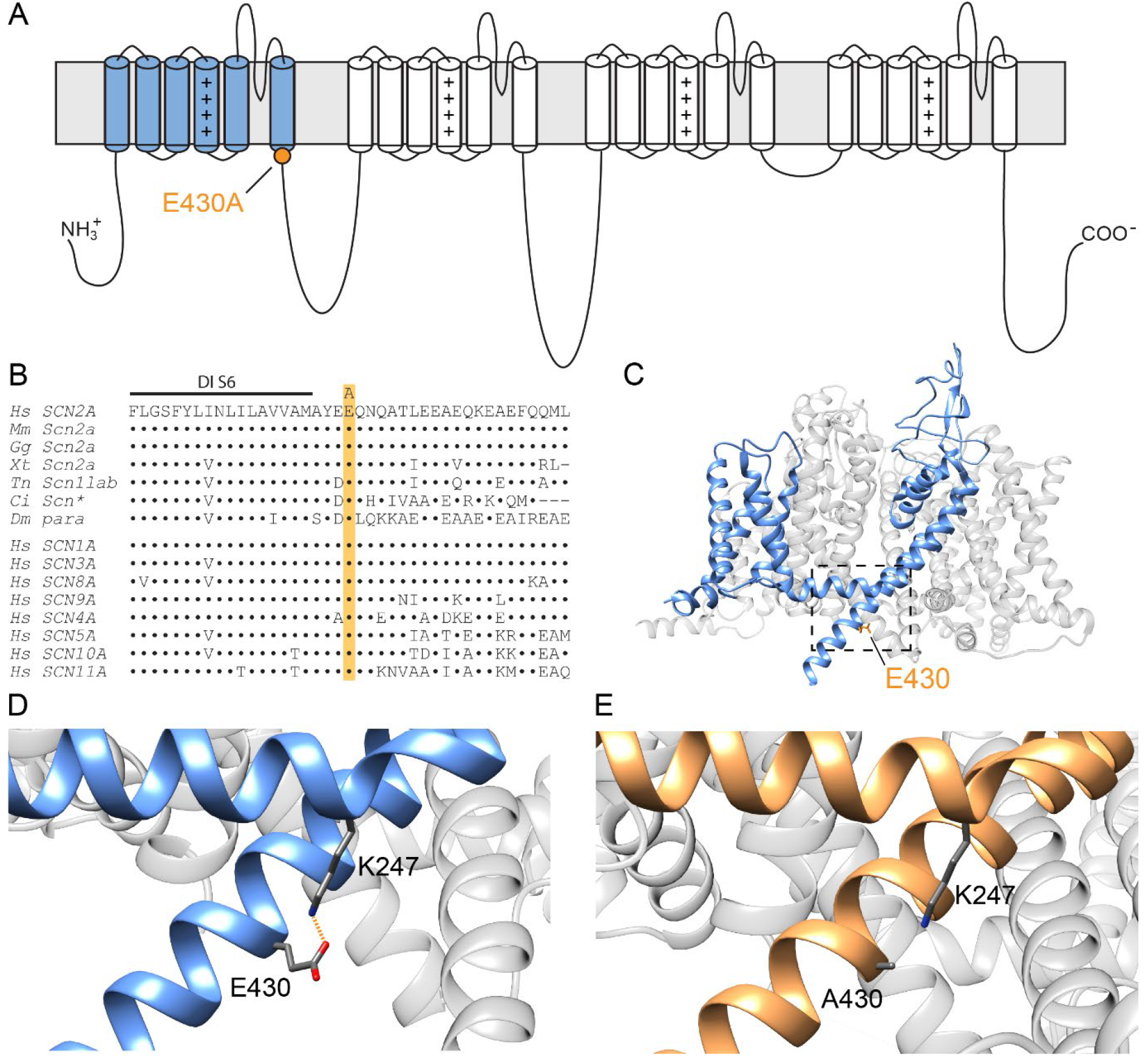
SCN2A-p.E430A location and predicted impact on Na_V_1.2. (**A)** Glutamate (E) at position 430 on Nav1.2 membrane topology map shows the location in the proximal DI-DII loop just beyond the D1S6 transmembrane segment. (**B**) Multiple alignment showing that E430 is invariant across SCN2A orthologs and paralogs. Hs, Homo sapiens; Mm, Mus musculus; Gg, Gallus gallus; Xt, Xenopus tropicalis; Tn, Tetraodon nigrovidis; Ci, Ciona intestinalis; Dm, Drosophila melanogaster. (**C)** (PDB: 6J8E, beta subunit and non-protein molecules omitted for clarity) with domain 1 shown in blue and E430 side chain marked in orange. The dotted box focuses on the S4-5 linker and proximal DI-DII cytoplasmic loop as shown in panels C and D. (C) Zoom in of the wild type AlphaFold3 structure highlighting the interaction between labeled E430 and K247. (D) Corresponding SCN2A-p.E430A AlphaFold3 structure with domain 1 colored in orange showing the loss of interaction due to alanine substitution.

We performed whole-cell voltage clamp recordings in HEK293T cells to assess the functional properties of wild-type (WT) Na_V_1.2 or E430A. While whole-cell current amplitude was similar between WT and E430A channels, the voltage-dependence of activation was significantly hyperpolarized for E430A channels (WT:-17.7 ± 1.0mV, n = 16, E430A:-22.2 ± 1.2mV, n = 17, p = 0.0088), (**Fig. 2A,B**, **Table 2**). Interestingly, we observed no difference in any other functional property that we assessed, including voltage-dependence of inactivation, recovery from inactivation, frequency-dependent rundown, or persistent sodium current. (**Fig. 2**, **Table 2**).

**Figure 2.**
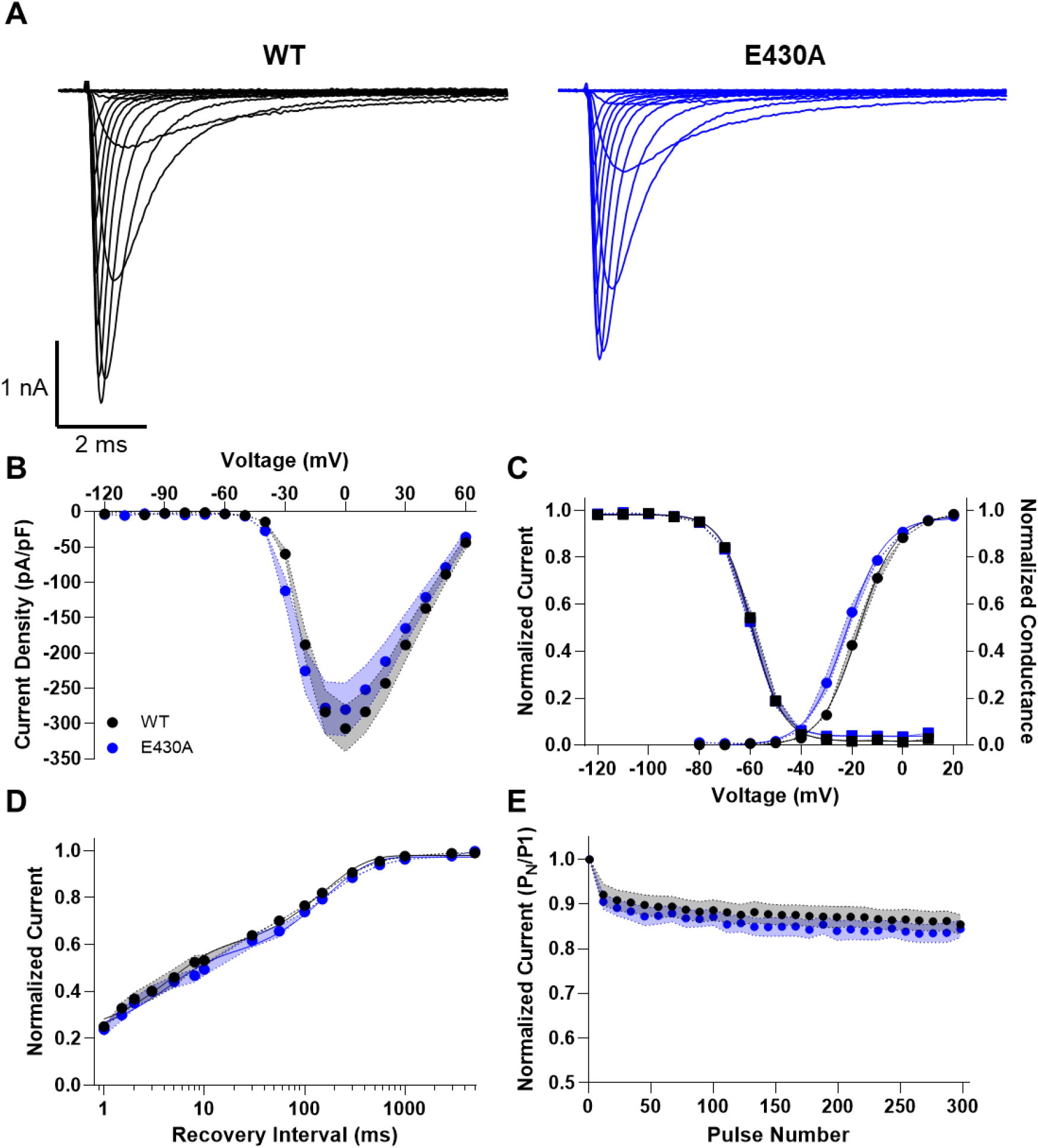
Heterologous expression of Nav1.2-E430A in HEK39T cells reveals altered voltage-dependence of activation. (**A**). Average whole-cell sodium currents for WT (left) and E430A (right) Na_V_1.2 channels. (**B**). Summary current-voltage relationship for WT (black) and E430A (blue) expressing cells. (**C**). Summary voltage-dependence of activation (circles) and inactivation (square) for WT (black) and E430A (blue) expressing cells. (**D**). Recovery from inactivation for WT (black) and E430A (blue) expressing cells. (**E**) Frequency-dependent rundown at 50 Hz for WT (black) and E430A (blue) expressing cells. All data are plotted and mean ± SEM for n = 16-17 cells.

### 3.2 Generation and initial characterization of Scn2a^E430A^ mice

The SCN2A-p.E430A variant was introduced into mouse *Scn2a* using CRISPR/Cas9 editing and homology directed repair in B6 embryos at the two-cell stage (**Fig. 3A**). Heterozygous *Scn2a^E430A^* (*Scn2a*^A/+^) and WT mice were born at the expected Mendelian ratios and had no survival deficit. In contrast, homozygous *Scn2a*^A/A^ mice exhibited perinatal mortality, consistent with reports from other *Scn2a* mutant mouse models (Echevarria-Cooper et al., 2022; Planells-Cases et al., 2000). Accordingly, all subsequent experiments focused on *Scn2a*^A/+^ mice, modeling the heterozygosity of individuals with *SCN2A*-related disorders. Forebrain expression of *Scn2a* transcript and Na_V_1.2 protein levels did not differ between *Scn2a*^A/+^ and WT littermates (**Fig. 3B-D**). Together, these data indicated that heterozygosity for the *Scn2a*^A/+^ allele does not alter baseline *Scn2a* expression, supporting its use as a GoF model for *in vivo* and *ex vivo* studies.

**Figure 3.**
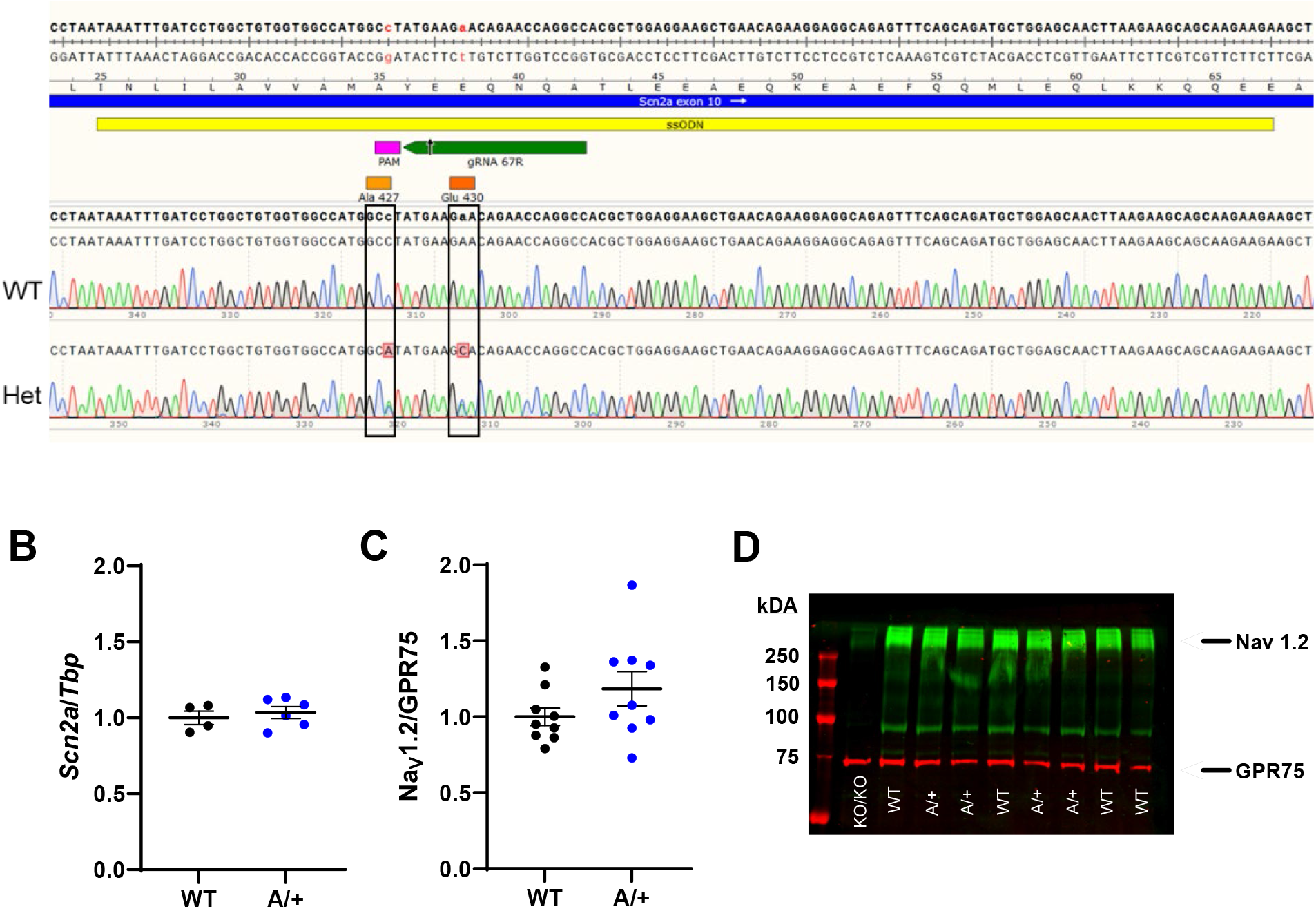
Generation and characterization of *Scn2a^E430A^* mice. (**A)** Schematic of CRISPR/Cas9 targeting strategy. (**B)** *Scn2a* transcript expression normalized to *Tbp* did not differ between WT and *Scn2a*^A/+^ mice (WT: 1.00 ± 0.04; A/+: 1.04 ± 0.09; p>0.57, n = 4-6/genotype). (**C)** Na_V_1.2 protein expression normalized to GPR75 did not differ between WT and Scn2a^A/+^ mice (WT: 1.00 ± 0.06; A/+: 1.19 ±0.11; p>0.16, n = 9/genotype). (**D)** Representative immunoblot for Na_V_1.2 (∼260 kDA, green) and GPR75 (∼72 kDA, red). A homozygous knockout (KO/KO) sample was included as a negative control. In B and C, symbols represent individual mice, horizontal lines indicate mean and error bars represent SEM.

### 3.3 Altered intrinsic excitability in Scn2a*^A/+^* hippocampal pyramidal neurons

We compared neuronal excitability from cultured hippocampal neurons (DIV14-16) from WT and *Scn2a^A/+^* mice. We observed that neurons from *Scn2a*^A/+^ mice were hyperexcitable compared to WT neurons (**Fig. 4**). Additionally, *Scn2a^A/+^*neurons showed a lower rheobase current, a hyperpolarized action potential threshold, and a larger fast AHP amplitude compared to WT neurons (**Fig. 4D,E,G**, **Table 2**). Other intrinsic neuronal properties, including resting membrane potential, action potential amplitude, half-width, upstroke and downstroke velocity, and input resistance were unchanged between WT and *Scn2a*^A/+^ neurons (**Fig. 4C,F,H-J**, **Table 2**).

**Figure 4.**
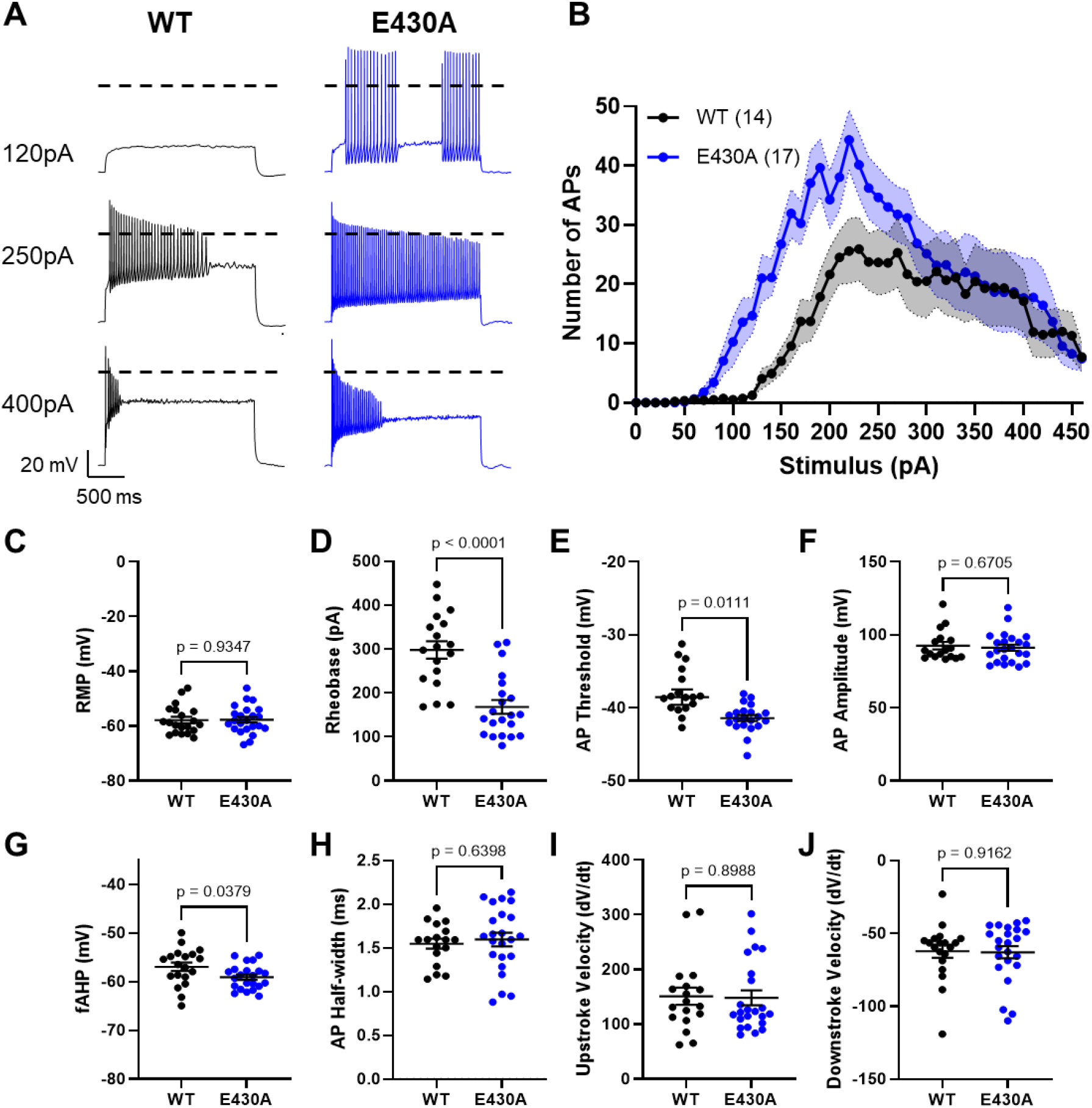
Altered excitability of hippocampal pyramidal neurons from *Scn2a*^A/+^ mice. **(A)** Representative action potential traces elicited by 120, 240 and 400 pA current injections for WT and E430A neurons. **(B)** Input-output curves for WT (black) and E430A (blue) neurons. **(C)** Resting membrane potential for WT (black) and E430A (blue) neurons. **(D)** Rheobase for WT and E430A neurons. **(E)** Action potential threshold for WT (black) and E430A (blue) neurons **(F)** Action potential amplitude for WT (black) and E430A (blue) neurons. **(G)** Fast afterhyperpolarization (fAHP) for WT (black) and E430A (blue) neurons. **(H)** Action potential half-width for WT (black) and E430A (blue) neurons. **(I)** Upstroke velocity for WT (black) and E430A (blue) neurons. **(J)** Downstroke velocity for WT (black) and E430A (blue) neurons. All data are plotted as mean ± SEM for n = 19-24 cells.

### 3.4 Enhanced LTP in Scn2a*^A/+^* hippocampal slices

To investigate the impact of the gain-of-function variant on hippocampal circuit integration, we compared measures of short-term and long-term plasticity between genotypes. We first assessed presynaptic function in the Schaffer collateral-CA1 pathway (**Fig. 5A**) using a paired-pulse ratio (PPR) protocol across a range of interstimulus intervals (25, 50, 100, 200, and 500 ms). PPR was calculated as the slope of the second fEPSP (P2) divided by the slope of the first (P1). PPR did not differ significantly between *Scn2a*^A/+^ mutant mice and wildtype littermate controls (**Fig. 5B**, **Table 2**), indicating that basal presynaptic release probability at Schaffer collateral-CA1 synapses is unaffected in mutant mice. We next examined long-term potentiation (LTP) induced by theta-burst stimulation (TBS) of the Schaffer collateral pathway. As expected, the stimulation paradigm reliably evoked long-lasting LTP (up to 60 minutes), evidenced by a steeper slope of the fEPSP in CA1. Interestingly, *Scn2a*^A/+^ mice (n = 8 slices) exhibited significantly greater LTP magnitude compared to wildtype controls (n = 7 slices), quantified during the 50–60-minute post-stimulation window normalized to baseline (**Fig. 5D**)(A/+: 197.9 ± 22.8%; WT: 134.8 ± 7.06%; *p* < 0.03). These findings indicate that mice expressing the GoF variant E430A have increased hippocampal long-term potentiation relative to wildtype, while short-term synaptic release is unchanged.

**Figure 5.**
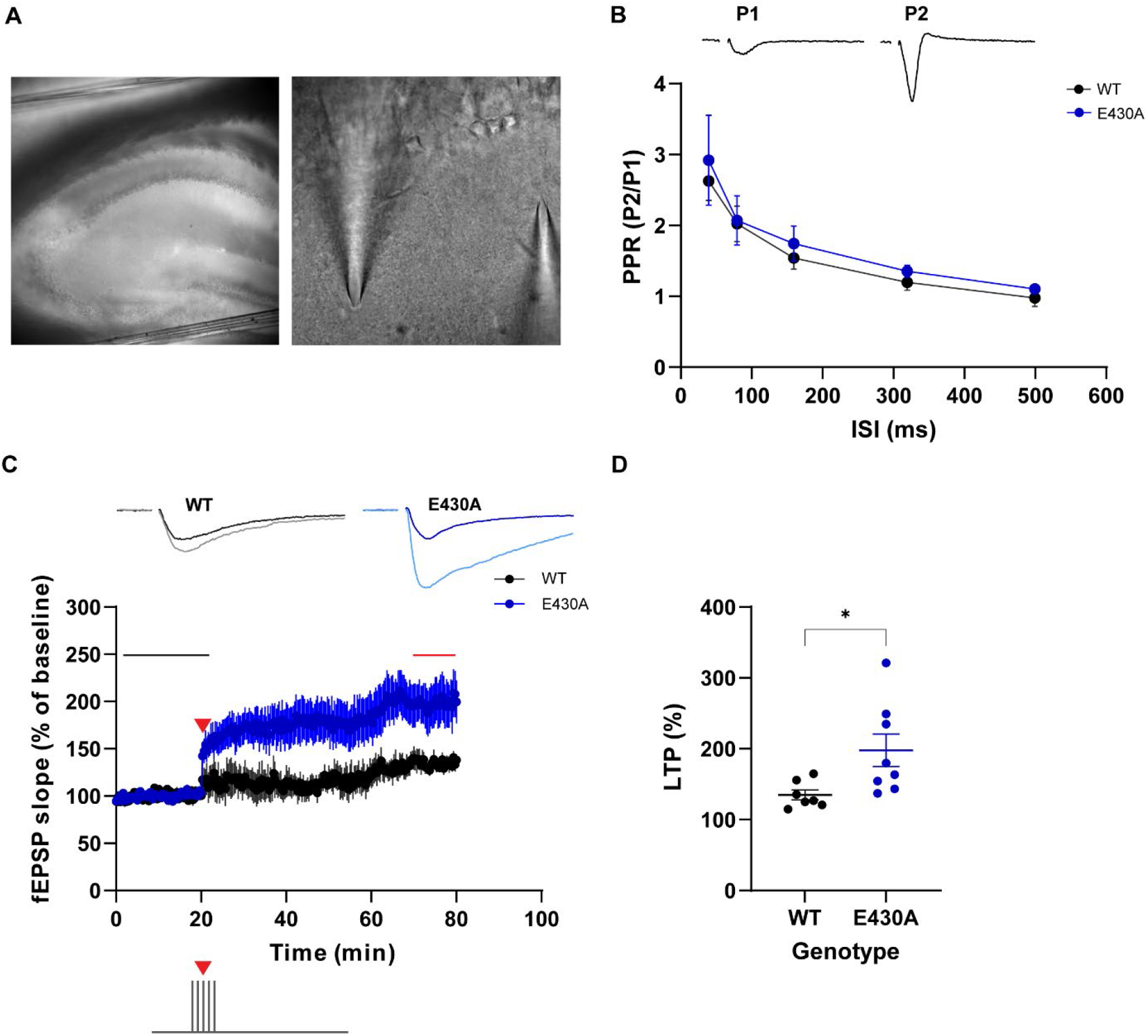
Hippocampal CA1 long-term potentiation is enhanced in *Scn2a*^A/+^ mice. **(A)** Representative images of hippocampal slice preparation and electrode placement for extracellular field recordings in area CA1 at low magnification (10x, left) and high magnification (40x, right). **(B)** Paired-pulse ratio across a range of interstimulus intervals (ISIs) in *Scn2a*^A/+^ mice (blue, n=6 slices) and littermate wildtype controls (black, n=6 slices). **(C)** Time course of field EPSP (fEPSP) slope (expressed as percentage of baseline) before and after theta burst stimulation protocol (TBS; red arrowhead, delivered after 20 minutes of baseline) in *Scn2a*^A/+^ (blue, n=8 slices) and W (black, n=7 slices). Representative fEPSP traces from each group are shown above at baseline and post-TBS. **(D)** Quantification of LTP magnitude expressed as mean fEPSP slope 50-60 minutes post-TBS normalized to baseline (gray bar on C). *Scn2a*^A/+^ mice displayed significantly greater LTP compared to WT controls (Welch’s t-test p=0.0288).

### 3.5 EEG abnormalities in Scn2a*^A/+^* mice

To assess baseline cortical activity, continuous video-EEG recordings were obtained from adult WT and *Scn2a*^A/+^ mice for at least 7 days, and the first two light and dark cycles were manually analyzed. Root mean square (RMS) did not differ by genotype during either the light or dark phases, indicating comparable overall EEG amplitude and baseline activity levels (**Fig. 6A**) (*Light*: WT: 19.1 ± 1.90 µV; A/+ 14.3 ± 2.44 µV; p > 0.13; *Dark*: WT 17.1 ± 2.48 µV; A/+ 15.4 ± 2.79 µV; *p* > 0.64).

**Figure 6.**
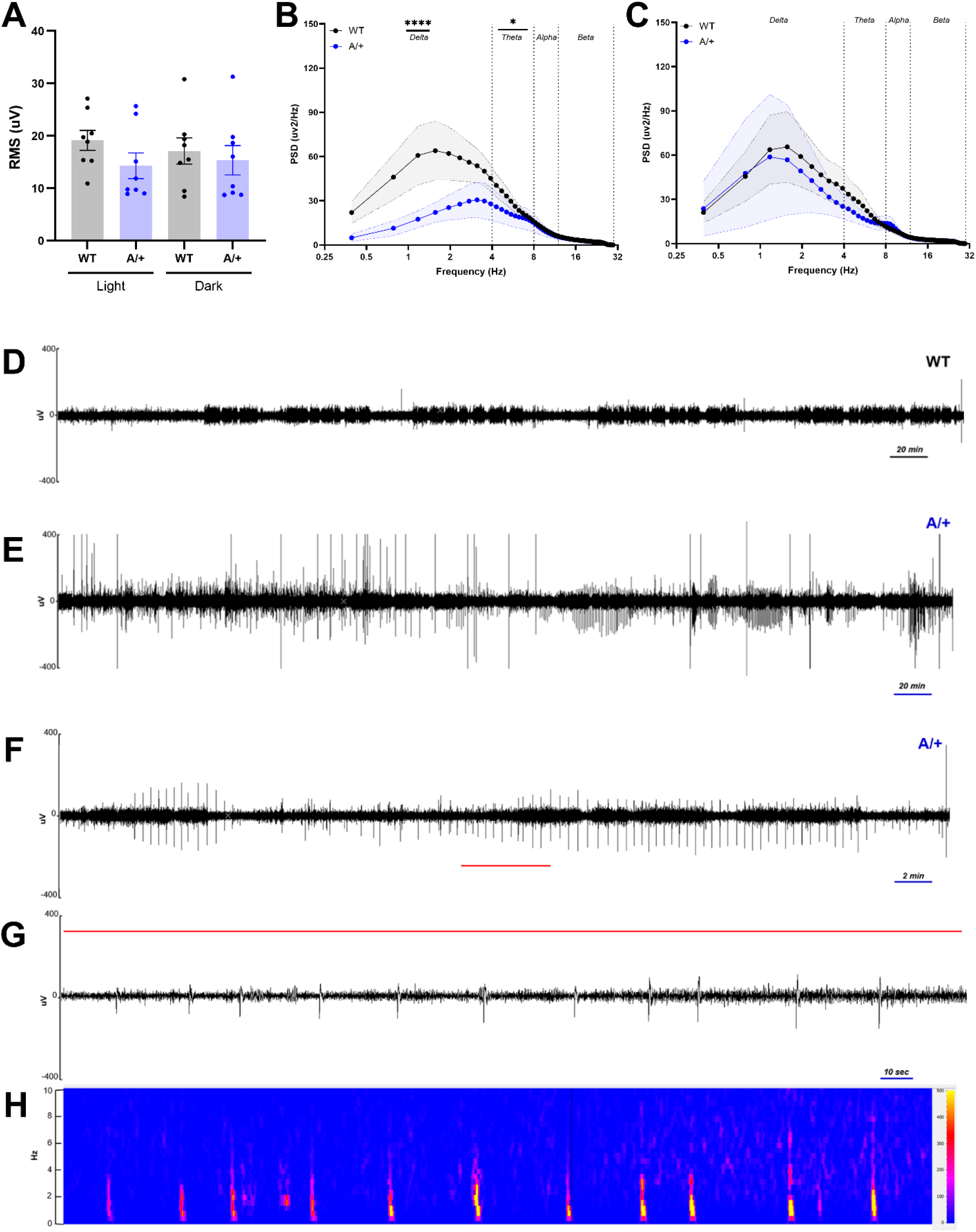
Baseline EEG characteristics of *Scn2a*^A/+^ (A/+) mice. **(A)** Root mean square (RMS) during light and dark cycles did not differ between genotypes (Light - WT: 19.1 ± 1.9 µV, A/+ 17.1 ± 2.5 µV; Dark - WT: 14.3 ± 2.4 µV; A/+: 15.4 ± 2.8 µV; n = 8 per genotype). Symbols represent individual mice for each light cycle phase, horizontal lines indicate mean and error bars represent SEM. **(B-C)** PSD during light (B) and dark (C) phases with frequency bands demarcated by dashed lines and shaded regions indicating SEM (n = 4 per genotype). During the light phase (B), *Scn2a*^A/+^ exhibited markedly lower reduction in delta (0-4 Hz, p<0.0001) and Theta (4-8 Hz, p = 0.0287) frequency bands compared to WT (n = 4 per genotype). (**D-E)** Representative ∼8-hour baseline EEG traces during light phase from a WT (D) or *Scn2a*^A/+^ mouse (E). The *Scn2a*^A/+^ EEG trace shows high-amplitude, sharp interictal epileptic spiking. (**F)** A ∼45 min. expanded view of green-lined section in E. **G)** A ∼5 min. expanded view of red-lined section in F. **H)** Time-frequency spectrogram corresponding to the EEG segment in G, demonstrating transient broadband power increases.

Despite similar RMS, power spectral density (PSD) analysis revealed genotype-dependent differences during the light phase. *Scn2a*^A/+^ mice exhibited a significant reduction in low-frequency power bands compared with WT mice, with decreased power in the delta (0-4 Hz; F(1,154) = 29.10, *p* < 0.0001) and theta (4-8 Hz; F(1,140) = 4.888, *p* = 0.0287) bands (**Fig.6B**). No differences were observed in alpha or beta frequency bands. In contrast, there were no genotype-dependent differences across any frequency band during the dark phase (**Fig. 6C**).

Manual inspection of EEG recordings revealed recurrent high-amplitude interictal epileptiform discharges in *Scn2a*^A/+^ mice compared to WT mice that had stable baseline activity (**Fig.6D-G**). Interictal events occurred as isolated discharges or brief trains (< 2 min) and were identified in all *Scn2a*^A/+^ mice. Spectral analysis of the interictal epileptiform discharges revealed transient broadband power increases (**Fig. 6H**). No spontaneous generalized seizures were detected in any of the recordings (176 of hours from 4 of mice).

### 3.6 Seizure susceptibility and phenytoin responsiveness in Scn2a^A/+^ mice

To evaluate seizure susceptibility, *Scn2a*^A/+^ and WT mice were assessed using three distinct seizure-induction paradigms, with or without phenytoin (PHT) pre-treatment. First, we assessed susceptibility to generalized seizures using ECS with 32 mA stimulation, which is subthreshold for B6 mice. *Scn2a*^A/+^ mice were more susceptible, with 80% (8/10) progressing to HLE followed by death, whereas no WT mice reached HLE (*p* < 0.0001)(**Fig. 7A**). Pretreatment with PHT completely prevented HLE in *Scn2a*^A/+^ mice (*p* < 0.0001), indicating robust protection against ECS-induced seizures, consistent with its known pharmacological profile in WT mice.

**Figure 7.**
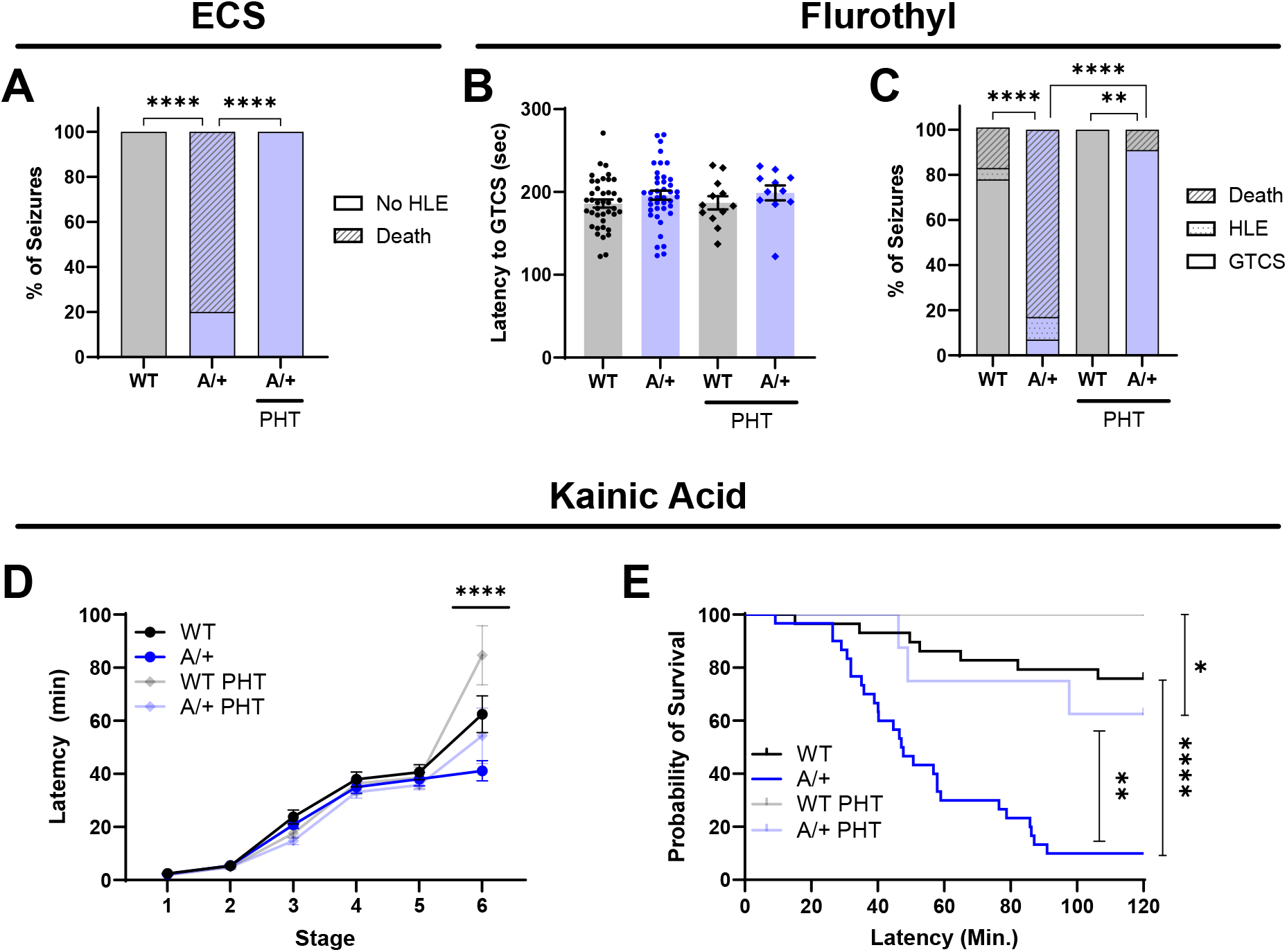
Seizure susceptibility and PHT sensitivity of *Scn2a^E430A^* mice. (**A)** *Scn2a*^A/+^ mice were more sensitive to a subthreshold electroconvulsive shock (ECS), with 80% (8/10) exhibiting HLE followed by death compared to no HLE in WT mice (p < 0.0001, n = 10/genotype). Pretreatment with PHT protected *Scn2a^A/+^* mice from HLE (p<0.0001, n=10/genotype). (**B)** Latency to GTCS following flurothyl exposure was unchanged by genotype or PHT treatment. GTCS latency was 196 ± 5.51 sec for *Scn2a^A/+^* mice and 186 ± 4.84 sec for WT. Following PHT pretreatment, *Scn2a^A/+^* exhibited GTCS latency of 199 ± 9.0 sec and WT GTCS latency was 187 ± 8.08 sec. (**C)** Seizure severity outcomes following flurothyl were affected by genotype. For *Scn2a*^A/+^ mice, 83% (34/41) exhibited lethality, 10% (4/41) HLE and 7% (3/41) GTCS only, compared to WT mice that had 18% (7/40) lethality, 5% (2/40) HLE and 78% (31/40) GTCS only (p < 0.0001). PHT pretreatment improved seizure outcomes for both genotypes, with 91% (10/11) of *Scn2a^A/+^* mice exhibiting GTCS only and just a single mortality (1/11) (p<0.0001), and 100% (12/12) of WT mice with GTCS only (p < 0.0001). Despite PHT protection for both groups, severity still differed between the genotypes (p = 0.0021). (**D)** Following KA administration, latency to reach the highest seizure stage (GTCS, stage 6), was affected by genotype. For *Scn2a*^A/+^ mice, 93% (28/30) reached GTCS with a latency of 41.2 ± 3.77 min compared to 59% (17/29) of WT mice with a latency of 62.4 ± 6.93 min (p<0.0001). PHT pretreatment did not significantly affect GTCS latency for *Scn2a*^A/+^ mice, with 63% (5/8) reaching GTCS with latency of 54.3 ± 10.5 min. In contrast, GTCS latency for WT mice was attenuated by PHT (84.7 ± 11.1 min; p<0.0001). The genotype-dependent difference in GTCS latency persisted with PHT pretreatment (p<0.0001). (**E)** Survival post KA administration was affected by genotype, with 90% (27/30) lethality for *Scn2a*^A/+^ mice compared to 24% (7/29) of WT (p < 0.0001). PHT pretreatment improved survival for *Scn2a*^A/+^ mice, with 38% (3/8) lethality (p < 0.005), although it did not normalize to the complete survival observed for WT mice (13/13) (p < 0.02).

Next, we evaluated generalized seizure susceptibility using the GABAergic antagonist flurothyl. Latency to GTCS following flurothyl exposure was not affected by genotype or by PHT pretreatment (**Fig. 7B**). However, seizure severity outcomes differed between genotypes (p < 0.0001)(**Fig. 7C**). The majority of *Scn2a*^A/+^ mice (93%) progressed to severe outcomes, including HLE (4/41) or HLE followed by death (34/41), while only 22% of WT mice progressed to severe outcomes (9/40). PHT pretreatment prevented severe outcomes in both genotypes, with only a single mortality for *Scn2a*^A/+^ mice and none for WT mice (*p* < 0.004) (**Fig. 7C**).

Finally, we evaluated seizure susceptibility using the L-glutamic acid analog kainic acid (KA) that induces slower progression through seizure stages, enabling finer resolution of seizure progression. Relative to WT mice, *Scn2a*^A/+^ mice followed a similar progression through less severe seizure stages but were more likely to reach the most severe stage of GTCS (stage 6) with shorter latency (**Fig. 7D**). Among *Scn2a*^A/+^ mice, 93% (28/30) of mice reached GTCS with a latency of 41.2 ± 3.77 min, while 59% (17/29) of WT reached GTCS with a latency of 62.4 ± 6.93 min (*p*<0.0001). Furthermore, *Scn2a*^A/+^ mice had higher rates of death following KA administration, with 90% (27/30) mortality in *Scn2a*^A/+^ mice compared to only 24% (7/29) in WT mice (*p* < 0.0001; **Fig 7E**). Pretreatment with PHT improved post-KA survival for *Scn2a*^A/+^ mice, with mortality of 38% (3/8); however, progression to GTCS was unaffected with 63% (5/8) of *Scn2a*^A/+^ mice exhibiting GTCS and a latency of 54.3 ± 10.5 min. Conversely, pretreatment of WT mice with PHT had no effect on survival but delayed progression to GTCS, with 31% (4/13) of WT mice reaching GTCS with a latency of 84.7 ± 11.1 min (**Fig. 7E**). Genotype-dependent differences in progression to GTCS and survival persisted for PHT pretreatment groups.

Thus, with multiple induction paradigms, *Scn2a^A/+^* mice show similar latency to generalized seizures, but exhibit enhanced evolution and poor outcomes.

## 4. Discussion

In this study, we generated and characterized a mouse model carrying the patient-associated *SCN2A*-p.E430A variant to investigate how selective alterations in activation gating influence neuronal excitability, circuit integration, and *in vivo* brain phenotypes. We demonstrate that the E430A variant produces a gain-of-function phenotype resulting in neuronal hyperexcitability, enhanced hippocampal LTP at Schaffer collateral-CA1 synapses, altered brain activity with interictal epileptiform discharges, and enhanced sensitivity to multiple seizure inducers that can be attenuated by the anti-seizure medication phenytoin. Together, these findings establish *Scn2a*^A/+^ mice as a clinically relevant and pharmacologically tractable model of *SCN2A*-related DEE.

The *Scn2a*-E430A variant suggests a structural explanation for the gain of function phenotype. E430 is located near the D1 S4-S5 linker, a region involved in coupling voltage-sensor movement to channel opening (Catterall, 2012). Structural modeling predicts the E430A variant disrupts a hydrogen bond with K247, potentially lowering the threshold for channel opening. This ability to open more readily or frequently could account for the hyperpolarized shift in activation. While this prediction is based on modeling, the cellular data presented here is consistent, with Nav1.2 E430A channels exhibiting a hyperpolarized shift in the voltage-dependence of activation without other biophysical alterations.

The GoF effect at the channel level results in hyperexcitability at the neuron level, with lower rheobase and AP threshold and larger fast AHP observed in excitatory pyramidal neurons isolated from *Scn2a*^A/+^ mice. Further, our findings of greater LTP magnitude in the Schaffer collateral-CA1 hippocampal circuit indicate that the E430A mutation enhances activity-dependent synaptic strengthening in hippocampal CA1 without altering release probability. This points to a post-synaptic or network-level mechanism underlying the augmented, likely aberrant plasticity (Marin-Castañeda et al., 2024). Conversely, mice with *Scn2a* haploinsufficiency, recapitulating LoF variants, have been shown to have impaired LTP (Shin et al., 2019; Spratt et al., 2019; Wang et al., 2024). These data further support the importance of SCN2A function in dendrites and suggest that disruption of SCN2A either by GoF or LoF results in dysregulation of LTP that may contribute to cognitive dysfunction.

The observed hyperexcitable state would support recurrent excitation and lower the barrier for interictal spikes and epileptiform activity (Bean, 2007; Jaffe and Brenner, 2018). Consistent with this, EEG recordings from *Scn2a*^A/+^ mice revealed recurrent interictal epileptiform discharges, with correlating spectral analysis showing transient broadband power increases in the absence of spontaneous seizure activity. Interictal epileptiform discharges are a recognized feature of *SCN2A*-related DEEs, reflecting latent network hyperexcitability and abnormal cortical synchronization that does not necessarily progress to spontaneous seizures (Sanders et al., 2018; Staley and Dudek, 2006; Wolff et al., 2017). Notably, reductions in low-frequency power were observed during the light phase, suggesting these effects are modulated by behavioral or arousal state, as previously described in both human and mouse epilepsy and animal models (Halász and Szűcs, 2020; Sevak et al., 2022).

Despite the absence of spontaneous seizures in EEG monitoring, *Scn2a*^A/+^ mice exhibited enhanced severity and lethality in response to proconvulsant stimuli. In the ECS assay, a stimulus that was subthreshold for WT mice resulted in near universal HLE and death in *Scn2a*^A/+^ mice. Similarly, *Scn2a*^A/+^ mice had high mortality following exposure to flurothyl, and accelerated progression to severe endpoints and near complete mortality following KA administration. Despite the enhanced severity and high mortality in chemoconvulsant assays, latency to initial seizure signs was not affected suggesting that the *Scn2a*-p.E430A GoF variant primarily affects seizure propagation or cessation rather than initiation. Consistent with this, prior work showed that *Scn2a* heterozygous deletion (LoF) resulted in shorter seizure duration and lower mortality in the *Kcna1*^-/-^ SUDEP model, while not affecting spontaneous seizure incidence or seizure threshold in response to flurothyl (Mishra et al., 2017). Together, these findings align with prior work demonstrating that network hyperexcitability can preferentially enhance seizure propagation and severity without altering initiation thresholds (Kriscenski-Perry et al., 2002; Quraishi et al., 2020; Santoro et al., 2010).

Phenytoin was reported to be effective for controlling neonatal-onset seizures in an individual with the *SCN2A*-p.E430A variant (Wolff et al., 2017). Furthermore, *SCN2A*-related DEEs with seizure onset before 3 months of age generally have good response to sodium channel blockers, particularly high-dose phenytoin (Howell et al., 2015; Sanders et al., 2018; Wolff et al., 2019; Wolff et al., 2017). Phenytoin treatment improved seizure outcomes across all induction paradigms, with particularly robust protection in the ECS and flurothyl models and partial protection in KA experiments. This pattern is consistent with the established efficacy of phenytoin, where PHT reliably limits seizure generalization and severity (Kriscenski-Perry et al., 2002; Putnam and Merritt, 1937; Swinyard et al., 1952). Importantly, the responsiveness of *Scn2a*^A/+^ mice supports the interpretation that this mutation confers a GoF sodium channel phenotype amenable to sodium channel blockers.

Although *Scn2a*^A/+^ mice did not exhibit spontaneous seizures as adults, it is possible they could have a transient seizure phenotype as was recently reported for *Scn2a^A263V^* heterozygotes (Reva et al., 2025). Future studies could examine early postnatal timepoints to determine if a similar self-limited seizure phenotype is evident. In addition, we and others have shown that genetic background can dramatically influence epilepsy phenotypes in mouse genetic models and crossing *Scn2a*^E430A^ mice with other inbred strains could unmask a more robust epilepsy phenotype (Bergren et al., 2005; Echevarria-Cooper et al., 2023; Hawkins et al., 2024; Kang et al., 2019; Löscher et al., 2017; Miller et al., 2014; Reva et al., 2025; Scott et al., 2025; Yu et al., 2020). The current study focused on electrophysiology and in vivo phenotypes related to seizures. Future studies will examine non-seizure phenotypes, including a comprehensive battery of neurobehaviors, particularly tests for hippocampal-dependent learning and memory and cognitive flexibility given our observation of enhanced hippocampal LTP.

## 5. Conclusions

In summary, our findings demonstrate that selective disruption of *Scn2a* activation gating is sufficient to produce neuronal hyperexcitability, interictal EEG abnormalities, heightened seizure severity, and sodium channel blocker responsiveness. The *Scn2a*^A/+^ mouse model captures key features of *SCN2A*-related DEE and provides a valuable platform for dissecting variant-specific mechanisms and evaluating targeted therapeutic strategies.

## Acknowledgements

We thank Conor Dixon, Talah Goldfarb, Rylie Pancoast, Nathan Speakes and Tyler Thenstedt for technical assistance. The genetically engineered mice were generated with the assistance of the Northwestern University Transgenic and Targeted Mutagenesis Laboratory. This work was supported by NIH grants R21-OD025330 (JAK), U54-NS108874 (ALG, JAK) and K08-NS121601 (SKA). Molecular graphics and analyses performed with UCSF Chimera, developed by the Resource for Biocomputing, Visualization, and Informatics at the University of California, San Francisco, with support from NIH P41-GM103311.

## CRediT authorship contribution statement

Conceptualization: NAH, CHT, SKA, ALG, JAK,

Data curation: NAH, CHT, DJIR

Formal analysis: NAH, CHT, DJIR, ET, SKA, JAK

Funding acquisition: ALG, JAK,

Investigation: NAH, DMEC, EEC, CHT, DJIR, ET

Methodology: NAH, DMEC, CHT, DJIR, ET, SKA, JAK

Project administration: NAH, JAK

Supervision: SKA, ALG, JAK

Visualization: NAH, CHT, DJIR, ET, SKA, JAK

Writing – original draft: NAH, DMEC, JAK

Writing – review & editing: NAH, DMEC, EEC, CHT, DJIR, ET, SKA, ALG, JAK,

## Materials and Data Availability

*C57BL/6J.Scn2a^em2Kea/Mmucd^* mice are available from Mutant Mouse Resources & Research Centers (MMRRC stock # 075779-UCD). Human Nav1.2 WT plasmid is available from Addgene (#162279). Datasets are available from https://doi.org/10.18131/qd5ca-dg744.

